# Educational attainment and personality are genetically intertwined

**DOI:** 10.1101/078014

**Authors:** René Mõttus, Anu Realo, Uku Vainik, Jüri Allik, Tõnu Esko

## Abstract

Heritable variance in psychological traits may reflect genetic and biological processes that are not necessarily specific to these particular traits but pertain to a broader range of phenotypes. We tested the possibility that Five-Factor Model personality domains and their 30 facets, as rated by people themselves and their knowledgeable informants, reflect polygenic influences that have been previously associated with educational attainment. In a sample of over 3,000 adult Estonians, polygenic scores for educational attainment (EPS; interpretable as estimates of molecular genetic propensity for education) were correlated with various personality traits, particularly from the Neuroticism and Openness domains. The correlations of personality traits with phenotypic educational attainment closely mirrored their correlations with EPS. Moreover, EPS predicted an aggregate personality trait tailored to capture maximum amount of variance in educational attainment almost as strongly as it predicted the attainment itself. We discuss possible interpretations and implications of these findings.

---

Personality trait variance has a substantial genetic component (Vukasović & Bratko, 2015). However, the specific genetic variants responsible for this have largely remained elusive, possibly due to a highly polygenic nature of the traits (Chabris et al., 2013). That is, large numbers of single-nucleotide polymorphisms (SNPs) collectively explain from nearly zero to under 20% of variance in personality traits, but the effect of any one SNP is usually too small to be reliably detectable (Smith et al., 2016; van den Berg et al., 2016; Lo et al., 2017). The same tends to be true for other psychological phenotypes such as intelligence (Davies et al., 2015) or subjective well-being (Okbay, Baselmans, et al., 2016). Slightly more variance has been traced to specific SNPs for some arguably less-psychological phenotypes such as educational attainment (Okbay, Beauchamp, et al., 2016) and body mass index (Locke et al., 2015).

It has also been suggested that personality traits could be conceived of as mostly phenotypic phenomena with limited or even no genetic or biological architecture of their own (Turkheimer, Pettersson, & Horn, 2014). If so, their observed genetic variance may to some, or perhaps even large, extent reflect genetic influences that act broadly across the organism as a nonspecific “genetic pull” rather than contribute to some systems specifically responsible for what appear as personality traits (Turkheimer et al., 2014). The genetic and resultant biological underpinnings of personality traits should then be shared with those of other phenomena that phenotypically relate to these personality traits but fall outside the personality domain *per se* (Mõttus, Marioni, & Deary, 2017).

Here, we address this possibility by investigating whether phenotypic variability in personality traits is associated with polygenic propensity for educational attainment (highest educational level obtained; henceforth education). Polygenic propensity refers to the combined additive effect of a large number of common SNPs captured in DNA arrays (i.e., SNPs in which the less prevalent alleles are not too rare). Numerous phenotypes may share genetic influences with personality characteristics. We chose education because it is a broad behavioral phenotype that has a sizable heritable component (Colodro-Conde, Rijsdijk, Tornero-Gómez, Sánchez-Romera, & Ordoñana, 2015; Silventoinen, Krueger, Bouchard, Kaprio, & McGue, 2004), is phenotypically correlated with a range of personality traits (Damian, Shanahan, Trautwein, & Roberts, 2015) and yet is not part of how the traits are usually operationalized—as self- or informant-reported summaries of thinking, feeling and behaving. Also, education has been relatively well characterized in terms of its associated SNPs (Okbay, Beauchamp, et al., 2016).

Twin studies have revealed that the phenotypic correlations of several personality traits with children's and adolescents’ academic results can largely be accounted for by shared genetic influences (Hicks, Johnson, Iacono, & McGue, 2008; Rimfeld, Kovas, Dale, & Plomin, 2016). In addition to the additive influences of individual genetic variants, these estimates reflect non-additive dominance and epistatic effects due to interactions between and within genetic loci, effects of rare variants and person-environment correlations (Purcell, 2002), and they are possibly confounded with environmental effects that twins share (Vinkhuyzen et al., 2012). Likewise, polygenic propensity for adult education has been linked with childhood self-control and interpersonal skills (Belsky et al., 2016), and low adult Neuroticism (Okbay, Beauchamp et al., 2016). These findings directly point to a possible overlap in the genomic correlates of personality traits and education. However, neither of these two studies addressed the implications of the findings for the genetic etiology of personality.

It is not known, however, whether such polygenic correlations with education are specific to these three personality traits or generalize to a wider spectrum of traits such as the domains and facets of the Five-Factor Model (FFM). To the extent that genetic variance in both education and personality does reflect a more general genetic pull, one would expect education to have a wider range of polygenic correlations with individuals’ characteristic patterns of thinking, feeling and behaving. Specifically, polygenic correlations should then be particularly likely for personality traits that are phenotypically correlated with education.

Employing published meta-analytic associations (Okbay, Beauchamp, et al., 2016) between years of education and SNPs, we created polygenic scores for education (EPS) for 3,061 adult Estonians. We correlated EPS with the five FFM domains and their 30 facets, as well as with an aggregate personality trait that combined, with optimal weights, education-related facets of personality. The range of 30 personality traits allowed us to test the ubiquity of polygenic correlations across the spectrum of personality characteristics. We employed both self- and informant-rated personality traits, which allowed for generalize the findings across specific assessment methods.

## Methods

### Sample

The current sample is a subset of the Estonian Biobank cohort (approximately 52,000 individuals), a volunteer-based sample of the Estonian resident adult population (Leitsalu et al., 2014). The participants were recruited randomly by general practitioners (GPs), physicians, or other medical personnel in hospitals or private practices as well as in the recruitment offices of the Estonian Genome Centre of the University of Tartu (EGCUT). Each participant signed an informed consent form, went through a standardized health examination and donated a blood sample for DNA. From among 3,426 individuals for whom both personality (self- and/or informant-reports) and DNA data were available, we selected 3,061 individuals (1,821 women) who were at least 25 years old (mean age 49.54 years, standard deviation 15.49, maximum 91) and had thereby had a chance to complete higher education and obtain a post-graduate degree. Apart from a slight over-representation of females (59%), the sample was a fairly representative cross-section of the adult Estonian population. For example, 37% of participants had higher education, which is comparable to the population estimate (http://stats.oecd.org). Personality data was collected only for the latest recruits of the EGCTU as the questionnaire was integrated at the last phase of the study.

### Measures

#### Personality

All but 15 of the selected participants (i.e., *N* = 3,046) completed the Estonian version of the NEO Personality Inventory 3 (NEO PI-3; McCrae & Costa, 2010), which is a slightly modified version of the Estonian version of the Revised NEO Personality Inventory (Kallasmaa, Allik, Realo, & McCrae, 2000). The NEO PI-3 has 240 items that measure 30 personality facets, which are then grouped into the five FFM domains, each including six facets consisting of eight items. The items were answered on a five-point scale (0 = *false/strongly disagree* to 4 = *true/strongly agree*). Personality traits of 2,904 of the selected participants (including the 15 participants with missing self-reports) were rated by an informant, who was typically spouse/partner, parent/child or friend. For cross-rater correlations, see Mõttus and colleagues (2014).

#### Education

Education was based on self-reports and quantified on an eight-level scale: without any formal education (*N* = 6), lower basic (*N* = 31), basic (*N* = 207), secondary (*N* = 550), vocational secondary (*N* = 956), applied higher (*N* = 177), higher (*N* = 967) or post-graduate education (*N* = 167). For the purpose of the analyses, the variable was treated as if it was based on an interval scale. [We repeated all analyses with education converted into years of education as per Okbay, Beauchamp et al. (2016) and obtained nearly identical results.]

#### Education polygenic scores

Genotyping was completed using different Illumina platforms (HumanCNV370-Duo and Quad BeadChip, OmniExpress BeadChips, HumanCoreExome-11 and HumanCoreExome-10 BeadChips) and the genotype data were imputed using the 1000 Genomes Project reference panel [Phase I integrated variant set release (v3) in NCBI build 37 (hg19)]. Imputed genotype probabilities were converted into hard-called genotypes using default settings in PLINK 1.9 software (Chang et al., 2015). In short, if imputation info metric (e.g. prediction uncertainty) value < .90, the variant was coded as missing, otherwise the genotype with the highest probability was used. As further quality control measures, SNPs with a minor allele frequency < 0.01 and Hardy Weinberg Equilibrium *p*-value < .001 were omitted from analyses.

Generally speaking, polygenic scores aggregate the small effects of large numbers of SNPs on a phenotype. The effect size for each SNP's designated allele (typically the less prevalent one), found in an independent sample, is multiplied by the count (0, 1, 2) of the allele for a given individual in the target sample, with the sums of these products across all SNPs then constituting the individual's polygenic score. For the current study, the effect sizes for individual SNPs were taken from a meta-analysis that linked over 8,000,000 SNPs with years of formal schooling (Okbay, Beauchamp, et al., 2016). The authors of the meta-analysis removed the contributions of the Estonian Genome Centre data from their combined results (discovery sample plus replication sample) for the purpose of the current study, so that the meta-analytic *N* varied from 100,000 to 319,946 depending on SNP. The genotypes were linkage disequilibrium-pruned using clumping to retain SNPs in linkage equilibrium with an *r*^2^ < .25 within a 250 bp window. The clumping procedure was carried out based on the subsample of 1,377 participants who had been genotyped using HumanCoreExome platforms in such a way that SNPs with lowest *p*-values in relation to education (in the meta-analysis of Okbay, Beauchamp, et al., 2016) were retained as the index SNPs of the clumps. No *p*-value cutoff was used for retaining SNPs. As a result of these procedures, individuals’ EPS values were based on over 150,000 SNPs (specifically, on 323,818 to 337,334 alleles). Ten principal components representing possible population stratification were calculated based on the genotype data and EPS was residualized for these components, as well as for the numbers of alleles contributing to EPS. The scores were calculated using PLINK1.9.

### Analyses

We first correlated individual FFM domains and facets with both EPS and education, controlling for age and sex. The *p*-values for each type of correlation (e.g., EPS with 35 self-reported personality traits) were adjusted for false discovery rate (Benjamini & Hochberg, 1995). Additionally, in order to efficiently capture the possibly multi-facet associations between personality and education, we weighed 30 personality facets by their unique associations with education and then aggregated them into a single composite variable. This composite represented personality polyfacet scores for education (*education polyfacet score*s); this was essentially analogous to how EPS aggregated the (mostly very small) effects of individual SNPs on education. In order to calculate the weights for each facet, we used the least absolute shrinkage and selection operator (LASSO) regression (Tibshirani, 2011) with 50-fold cross-validation and a shrinkage parameter lambda that minimized cross-validated error. This method effectively dealt with multi-collinearity among facets as well as reduced biases due to over-fitting. By nature, the polyfacet scores captured as much variance in education as could collectively be predicted by the 30 facets, even those that had not been significantly correlated with education in the bi-variate analyses. The scores could therefore be conceived of as reflecting an education-specific personality trait. We then carried out exactly the same procedure for the EPS. Both polyfacet scores were residualized for age and sex. These procedures were carried out separately for self- and informant-ratings of personality facets.

## Results

Polygenic propensity for education, EPS, had a correlation of .18 [95% confidence intervals (CI): .14, .21] with its target phenotype, education (here and henceforth, for all correlations reported in the text *p* < .001, unless reported otherwise). The association did not appear perfectly linear across seven levels of education, with medium levels of education being rather similar in their EPS scores (the average for the six people with no formal education is not shown in Figure 1). However, the average difference between people with lower basic education and a post-graduate degree was substantial (0.84 standard deviation units). Table 1 shows the phenotypic associations of personality traits with both phenotypic education and EPS (confidence intervals are reported in Table S1 in the Supplementary Material).

**Figure 1.**
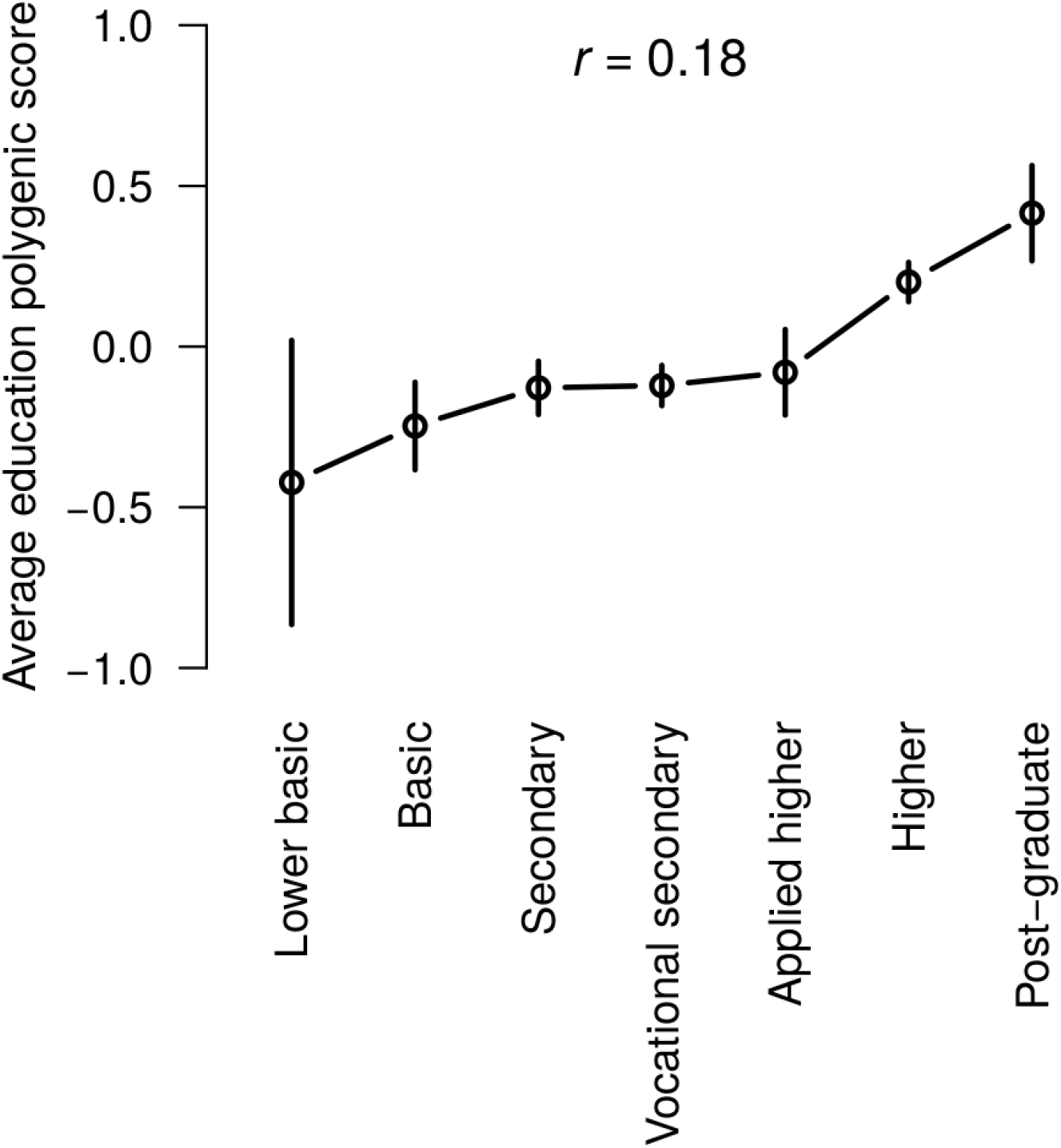
Educational levels and standardized education polygenic scores (EPS; means and 95% confidence intervals).

**Table 1.** Associations of personality domains and facets with education and education polygenic scores (EPS).

|  | Phenotypic education |  |  |  | EPS |  |  |  |
| --- | --- | --- | --- | --- | --- | --- | --- | --- |
|  | Self-ratings |  | Informants |  | Self-ratings |  | Informants |  |
|  | *r* | *p* | *r* | *p* | *r* | *p* | *r* | *p* |
| Neuroticism | **-.16** | <.001 | **-.17** | <.001 | **-.07** | <.001 | **-.05** | .022 |
| Extraversion | **.12** | <.001 | **.11** | <.001 | .02 | .248 | .01 | >.250 |
| Openness | **.25** | <.001 | **.19** | <.001 | **.11** | <.001 | **.09** | <.001 |
| Agreeableness | -.03 | .173 | .05 | .018 | -.01 | >.250 | .03 | .200 |
| Conscientiousness | **.07** | <.001 | **.13** | <.001 | -.04 | .074 | .02 | >.250 |
| N1: Anxiety | **-.11** | <.001 | **-.12** | <.001 | **-.06** | .004 | -.03 | >.250 |
| N2: Hostility | **-.17** | <.001 | **-.13** | <.001 | **-.08** | <.001 | **-.07** | .001 |
| N3: Depression | **-.17** | <.001 | **-.15** | <.001 | **-.07** | .001 | -.04 | .075 |
| N4: Self-Consciousness | **-.09** | <.001 | **-.12** | <.001 | **-.05** | .012 | -.02 | >.250 |
| N5: Impulsiveness | -.03 | .075 | **-.09** | <.001 | -.03 | .152 | -.04 | .079 |
| N6: Vulnerability to Stress | **-.14** | <.001 | **-.17** | <.001 | -.05 | .026 | -.03 | .232 |
| E1: Warmth | .04 | .044 | **.06** | .003 | .01 | >.250 | .01 | >.250 |
| E2: Gregariousness | **.05** | .003 | .04 | .023 | .00 | >.250 | .00 | >.250 |
| E3: Assertiveness | **.20** | <.001 | **.16** | <.001 | **.05** | .016 | .03 | .247 |
| E4: Activity | **.12** | <.001 | **.14** | <.001 | .01 | >.250 | .01 | >.250 |
| E5: Excitement Seeking | .03 | .189 | .03 | .124 | .01 | >.250 | -.01 | >.250 |
| E6: Positive Emotion | **.07** | <.001 | **.06** | .004 | .02 | >.250 | .01 | >.250 |
| O1: Openness to Fantasy | **.08** | <.001 | .04 | .052 | .04 | .029 | .03 | .199 |
| O2: Openness to Aesthetics | **.17** | <.001 | **.13** | <.001 | **.06** | .001 | **.06** | .007 |
| O3: Openness to Feelings | **.10** | <.001 | **.07** | <.001 | .03 | .216 | .00 | >.250 |
| O4: Openness to Actions | **.23** | <.001 | **.18** | <.001 | **.09** | <.001 | **.07** | .001 |
| O5: Openness to Ideas | **.24** | <.001 | **.24** | <.001 | **.12** | <.001 | **.11** | <.001 |
| O6: Openness to Values | **.20** | <.001 | **.09** | <.001 | **.11** | <.001 | **.06** | .004 |
| A1: Trust | **.22** | <.001 | **.16** | <.001 | **.09** | <.001 | **.06** | .010 |
| A2: Straightforwardness | .01 | >.250 | **.07** | <.001 | .03 | .128 | .03 | .179 |
| A3: Altruism | -.02 | >.250 | .03 | .115 | .00 | >.250 | .03 | >.250 |
| A4: Compliance | -.01 | >.250 | .03 | .118 | .01 | >.250 | .05 | .054 |
| A5: Modesty | **-.15** | <.001 | -.04 | .024 | **-.08** | <.001 | -.02 | >.250 |
| A6: Tendermindedness | **-.15** | <.001 | **-.06** | .001 | **-.09** | <.001 | -.02 | >.250 |
| C1: Competence | **.12** | <.001 | **.17** | <.001 | .01 | >.250 | .04 | .142 |
| C2: Order | .00 | >.250 | .01 | >.250 | **-.05** | .007 | -.03 | .190 |
| C3: Dutifulness | **.06** | .003 | **.11** | <.001 | -.04 | .033 | .03 | .199 |
| C4: Achievement Striving | **.06** | .001 | **.15** | <.001 | -.02 | .248 | .00 | >.250 |
| C5: Self-Discipline | .02 | >.250 | **.07** | <.001 | -.04 | .033 | .00 | >.250 |
| C6: Deliberation | **.08** | <.001 | **.13** | <.001 | .00 | >.250 | .04 | .079 |

NOTE: $r$ = correlations; $p$ = $p$ -values (adjusted for false discovery rate per column); informants = informant-ratings. Associations for which 99% confidence intervals did not span zero are marked in bold.

### Personality and polygenic propensity for education

In both self- and informant-ratings, EPS was significantly negatively correlated with the Neuroticism domain and positively correlated with the Openness domain, although the significance did not apply to all of their facets. Specifically, the associations were significant in both rating types for N2: Hostility, O2: Openness to Aesthetics, O4: Openness to Actions, O5: Openness to Ideas and O6: Openness to Values. The associations were also significant in both rating types for the A1: Trust facet of the Agreeableness domain. Some associations were only significant in self-reports; for example, people with higher EPS tended to rate themselves lower on A5: Modesty and A6: Tendermindedness, whereas this was not apparent in informant-ratings. In relative terms, the associations of EPS with self-reported facets were highly similar with its associations with informant-rated facets: the correlation between the two vectors of 30 correlations (Fisher-transformed) was .86 (CI: .73, .93).

What we find particularly noteworthy is that facets’ associations with EPS also closely mirrored their associations with phenotypic education. Specifically, facet-education and facet-EPS correlations (from Table 1, Fisher-transformed) strongly tracked each other in both self-reports [*r* = .91 (CI: .81, .96)] and informant ratings [*r* = .84 (CI: .69, .92)]. As shown in Figure 2, both associations were linear across the spectrum of effect sizes in that neither of them was driven by the few facets which significantly correlated with both education and EPS^1^. For example, even when we only considered facets which had an absolute correlation < .05 with EPS (i.e., mostly non-significant correlations), the facet-education and facet-EPS associations mirrored each other in both self-reports [*r* = .61 (CI: .21, .84), *p* = .007, *df* = 16] and informant-ratings [*r* = .78 (CI: .55, .90), *df* = 22].

**Figure 2.**
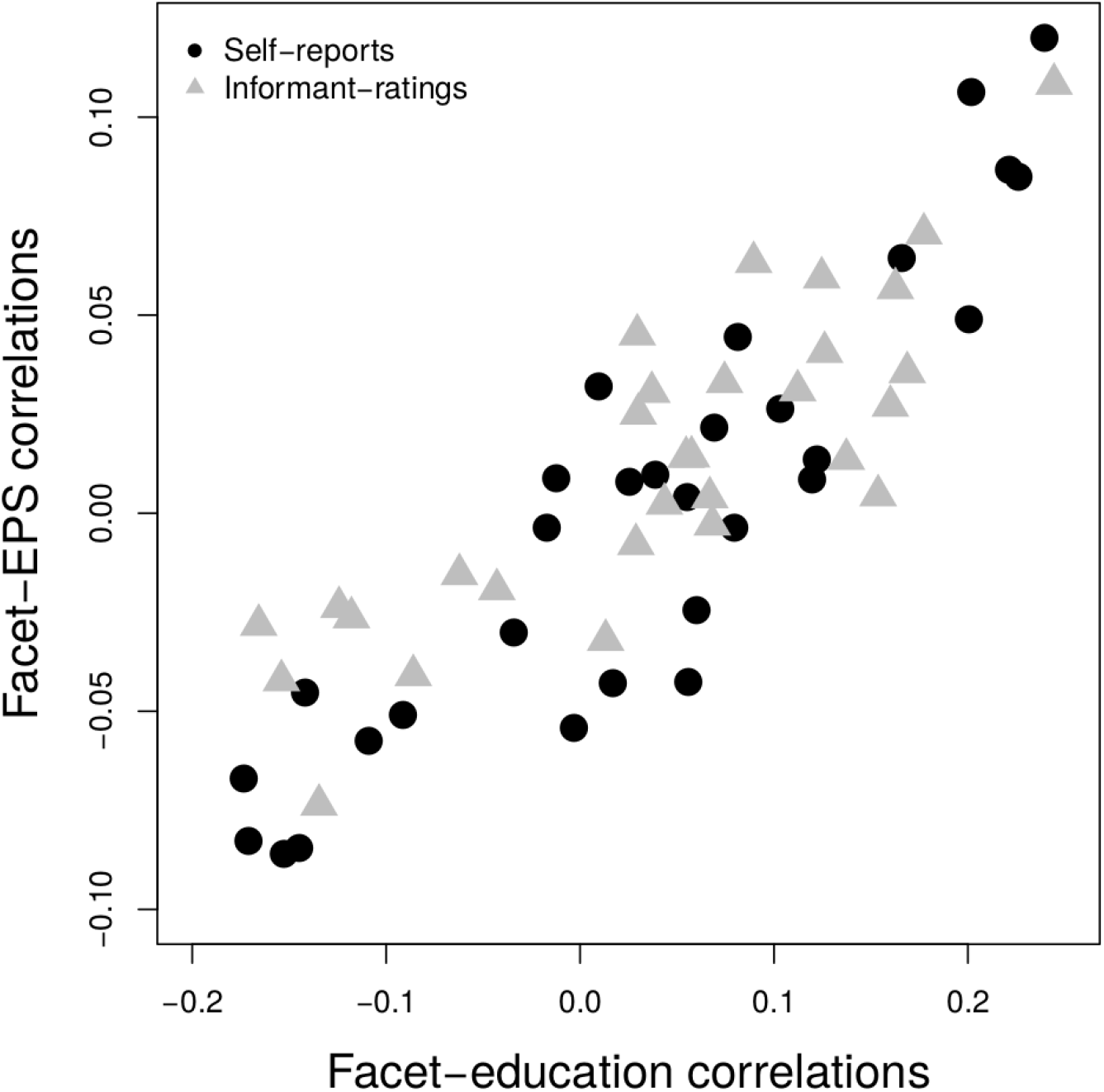
The associations of 30 personality facets with education and education polygenic scores (EPS).

### Associations with polyfacet scores

Education had sizable correlations (*r* = .39 to .45; Table 2) with its polyfacet scores. The correlations between education polyfacet scores and EPS were .17 and .14, respectively for self- and informant-ratings. These correlations suggest that the association of EPS with the education-related aspects of personality, appropriately aggregated, was nearly of the same magnitude than had been its correlation with the phenotypic education itself (the correlations were not significantly different with *p* > .05). Table 2 also provides partial correlations among the variables (i.e., correlations adjusting for the other two correlations) to gauge the extents to which either education polyfacet scores accounted for the effect of EPS on education or, conversely, education accounted for the effect of EPS on education polyfacet scores. The associations of EPS with both education and polyfacet scores were attenuated but remained substantially greater than zero, providing no clear evidence that either personality or education could (at least fully) mediate each other's polygenic influences. The correlations between the polyfacet scores for education and the similarly-created polyfacet scores for EPS were .81 (CI: .79, .82) and .79 (CI: .78, .81), respectively for self- and informant-ratings. These high correlations are consistent with Table 1 and Figure 2, showing that facet-education correlations closely mirrored facet-EPS correlations. All LASSO regression coefficients used for creating polyfacet scores are given in Table S2 (Supplementary Material).

**Table 2.** Zero-order and partial correlations between education, education polyfacet scores and education polygenic scores (EPS).

|  | Self-reported personality |  | Informant-reported personality |  |
| --- | --- | --- | --- | --- |
|  | *Zero-order correlations* |  |  |  |
|  | EPS | Education | EPS | Education |
| Education | .18 (.14, .21) |  | .18 (.14, .21) |  |
| Polyfacet scores | .17 (.14, .21) | .45 (.42, .48) | .14 (.11, .18) | .39 (.36, .42) |
|  | *Partial correlations* |  |  |  |
|  | EPS | Education | EPS | Education |
| Education | .12 (.08, .15) |  | .14 (.10, .17) |  |
| Polyfacet scores | .10 (.07, .14) | .43 (.40, .46) | .08 (.04, .12) | .37 (.34, .40) |

NOTE: 95% confidence intervals are given in brackets.

## Discussion

The results showed a systematic overlap between additive polygenic variance in education and personality. While previous research had reported polygenic correlations between education and a limited number of traits (Belsky et al., 2016; Okbay, Baselmans et al., 2016), we examined them across five FFM domains and their 30 facets, relying on one of the most comprehensive personality assessment frameworks currently available (McCrae & Costa, 2010). Education polygenic scores (EPS) correlated with several self- and informant-rated personality traits, especially those belonging to Neuroticism and Openness domains, and the associations closely mirrored the correlations of the traits with phenotypic education.

Although individual correlations between personality traits and EPS were small in absolute scale, they must be interpreted in the appropriate context. For example, polygenic scores for traits such as subjective well-being, depressive symptoms and Neuroticism account for less than 1% of variance in their respective traits (Okbay, Baselmans et al., 2016). This is similar to how polygenic scores for a *different* phenotype, education, predicted some personality traits in this study. Also, EPS was unlikely to capture full genetic variance in education and thereby also in related personality traits. For example, the heritability of education has been estimated at more than 20% based on alternative GWAS-derived procedures (Marioni et al., 2014), whereas EPS could only account for 3% of education phenotypic variance. Moreover, when we aggregated facets as per their association with education, the resulting correlations with EPS were comparable to its correlation with education itself.

There are multiple ways to interpret such polygenic overlap. One possible explanation is that the same genetic variants independently influence both education and personality, perhaps through some complicated biological and/or environmental pathways. Also, experiences related to education may be causal to personality traits and therefore genetic influences on education can account for some of the genetic variance in these traits. For example, certain genetic variants may predispose people to completing more years of schooling (e.g., via faster information processing or better physical health that allows for more engagement with education), which in turn may enhance people's interests in aesthetic and intellectual experiences or contribute to disapproval of dishonesty. In both cases, the genetic etiology of personality is at least partly entangled with that of education. Alternatively, personality traits may mediate the genetic variance in education (Rimfeld et al., 2016). Some traits may predispose people to seek out more schooling and their genetic influences can thereby account for some of the genetic variance in education, alongside any down-stream consequences of this important life-outcome such as job success or health. If so, the polygenic influences previously linked with education pertain, more proximally, to personality traits.

In order to assess the plausibility of these explanations, one could try to study people with no “exposure” to the hypothesized mediator (Kippersluis & Rietveld, 2016). If the otherwise observed polygenic correlations between education and personality traits are absent in people without formal education, this would support education being the mediating phenotype in these associations. However, applying the same logic to examine the mediating role of personality traits would be problematic because personality traits are never absent but only vary in degrees. Additionally, specific genetic variants with known causal pathways to the hypothesized mediator could be used to disentangle causality (Davey Smith, 2010), but too few, if any, genetic variants with clear causal pathways to personality traits and education are currently known.

The associations of the 30 personality facets with EPS closely mirrored their associations with education itself. This may provide indirect evidence against the possibility that the genetic effects captured by EPS *only* pertained to personality traits, which then phenotypically transmitted these effects to education. If this was the case, there would be no reason to expect the EPS-facet correlations to almost perfectly track education-facet association. Of course, this could happen due to unmeasured mediators—for example, biological or parental characteristics as well as other behavioral traits or life circumstances—linking EPS and education in addition to personality facets. We only controlled for age, sex and genetic stratification. But even then the genetic variance in personality would be entangled with that of education, with overlapping genetic variants in part independently contributing to both. However, EPS-facet associations mirroring education-facet associations are expected when both education and personality independently reflect the same genetic influences or when education mediates the genetic influences to personality.

Recently, Lo and colleagues (2017) provided evidence for sizable polygenic overlap between the FFM personality traits and a range of psychiatric phenotypes, as well as between the FFM traits themselves. There is also evidence for polygenic overlap between personality and some aspects of physical health such as body mass index and heart disease as well as health-relevant behaviors such as smoking (Gale et al., 2016). Importantly, most of these mental and physical health-related phenotypes also have polygenic correlations with education (Okbay, Beauchamp et al., 2016; Bulik-Sullivan et al., 2015). Combined with our results, this pattern of findings can be interpreted in the light of the hypothesis that the observed genetic variance in personality traits may at least partly reflect a general genetic pull—genetic influences that act broadly across a range of phenotypes rather than specifically on what have been operationalized as personality traits (Turkheimer et al., 2014).

Genetic variance of personality traits being partly entangled with that of education has implications beyond helping us to understand the etiology of the traits. First, attempts to delineate the specific genetic underpinnings of education or aspects of physical health may incidentally reveal the genetic mechanisms of phenotypically related personality traits. Also, these phenotypes could be used as proxies to narrow the range of potentially personality-related genetic variants as has been done for intelligence (Rietveld et al., 2014). Second, the genetic overlap needs to be factored into any attempts to interpret the phenotypic associations of personality traits with education and its associated characteristics such as those reflecting socioeconomic success. Turkheimer and colleagues (2014) argue that when associations of personality traits with other variables are investigated “our scientific hypotheses are usually phenotypic in nature” (p. 533). To the extent that genetic overlap is involved, there may be less of such phenotypic causation. Naturally, the implications of our findings stretch beyond the associations between personality traits and education. Genetic overlap should be considered for *any* phenomenon that is hypothesized to be either causal to behavioral traits or among their downstream consequences. For example, personality traits are phenotypically associated with obesity (Sutin, Ferrucci, Zonderman, & Terracciano, 2011), but these links may at least to some extent reflect genetic overlap. With genetic data becoming widely available, researchers will be increasingly able to decompose phenotypic associations into genetic and non-genetic components.

In sum, the current study examined polygenic overlap between education and a range of personality traits, and found clear evidence for this. There are various possible interpretations for this finding. In combination with recent evidence for genetic correlations between personality and various aspects of (mental) health, the regularity with which the genetic and phenotypic associations between personality traits and education mirrored each other suggests that genetic influences on personality may not necessarily pertain to some personality-specific neurobiological structures. Instead, genetic variance in personality traits may reflect the results of a more general genetic influences-related pull. Moreover, it is possible that this general pull extends to other psychological traits not addressed in this study such as attitudes, beliefs or motivation. Psychological phenomena are ubiquitously heritable, but they may not be aligned with distinct etiological mechanisms.

## Footnotes

1 These correlations could have been inflated by inter-facet differences in psychometric properties such as reliability or validity. However, even the differences between self- and informant-reports in facet-level correlations with EPS on one hand and education on the other mirrored each other [*r* = .80 (CI: .69, .90)]. This suggests that systematic (across facets) measurement inaccuracies were not a likely cause for the general similarity of the personality-genotype and personality-phenotype associations.

